# Hippocampal transcriptomic responses to cellular dissociation

**DOI:** 10.1101/153585

**Authors:** Rayna M. Harris, Hsin-Yi Kao, Juan Marcos Alarcón, Hans A. Hofmann, André A. Fenton

## Abstract

Single-neuron gene expression studies may be especially important for understanding nervous system structure and function because of the neuron-specific functionality and plasticity that defines functional neural circuits. Cellular dissociation is a prerequisite technical manipulation for single-cell and single cell-population studies, but the extent to which the cellular dissociation process affects neural gene expression has not been determined. This information is necessary for interpreting the results of experimental manipulations that affect neural function such as learning and memory. The goal of this research was to determine the impact of chemical cell dissociation on brain transcriptomes. We compared gene expression of microdissected samples from the dentate gyrus (DG), CA3, and CA1 subfields of the mouse hippocampus either prepared by a standard tissue homogenization protocol or subjected to a chemical cellular dissociation procedure. We report that compared to homogenization, chemical cellular dissociation alters about 350 genes or 2% of the hippocampal transcriptome. While only a few genes canonically implicated in long-term potentiation (LTP) and fear memory change expression levels in response to the dissociation procedure, these data indicate that sample preparation can affect gene expression profiles, which might confound interpretation of results depending on the research question. This study is important for the investigation of any complex tissues as research effort moves from subfield level analysis to single cell analysis of gene expression.

---

Nervous systems are comprised of diverse cell types that express different genes to serve distinct functions. Even within anatomically-defined subfields of the brain, there are identifiable sub-classes of neurons that belong to distinct functional circuits (***Danielson et al., 2016; Mizuseki et al., 2011; Namburi et al., 2015***). Cellular diversity is even greater when we consider that specific cells within a functional class can be selectively altered by neural activity in the recent or distant past (***Denny et al., 2014; Garner et al., 2012; Ramirez et al., 2013; Reijmers et al., 2007***). This complexity can confound the interpretation of transcriptome data collected from bulk samples containing hundreds to tens of thousands of cells that represent numerous cellular subclasses at different levels of diversity.

Recent advances in tissue harvesting and sequencing technologies have allowed detailed analyses of genome-scale gene expression profiles at the level of single-cell populations in the context of brain and behavior studies (***Mo et al., 2015; Chalancon et al., 2012; Lacar et al., 2016; Moffitt et al., 2018; Nowakowski et al., 2018; Raj et al., 2018***). These approaches have led to systems-level insights into the molecular substrates of neural function and to the discovery and validation of candidate pathways regulating physiology and behavior. Current methods for dissociating tissues into single-cell suspensions include mechanical and enzymatic treatments (***Jager et al., 2016***). To complement the efforts allowing for single-neuron analysis of transcriptional activity, it is necessary to understand the extent to which the dissociation treatment of tissue samples prior to single-cell transcriptome analysis might confound interpretation of the results.

Our experiment was designed to determine if enzymatic dissociation itself alters the transcriptome of the hippocampus. We did not compare single-cell RNA-seq data to bulk tissue RNA-seq data because that is orthogonal to the present research question. Instead, we compare transcriptome data from the CA1, CA3, and dentate gyrus (DG) subfields of the hippocampus subjected to one of two treatments 1) homogenized (HOMO) or 2) dissociated (DISS). Samples were prepared by a standard homogenization protocol and the sequencing results were compared to corresponding samples that were dissociated as if they were being prepared for single-cell sequencing (***Figure 1***A). Importantly, the dissociated tissue was not sorted or differentially treated in any way further, which would of course defeat the purpose of dissociation for single cell or single cell population studies, but is essential for the task at hand. Accordingly, we could expect the same tissue constituents in the two groups, and can therefore attribute differences in gene expression to the treatment procedure. We used the Illumina HiSeq platform for sequencing, Kallisto for transcript abundance estimation (***Bray et al., 2016***) and DESeq2 for differential gene expression profiling (***Love et al., 2014***). Data and code are available at NCBI’s Gene Expression Omnibus Database (accession number GSE99765), as well as on GitHub (https://github.com/raynamharris/DissociationTest) with an archived version at the time of publication available on Zenodo (***Harris, 2019***). A more detailed description of the methods is provided in the supplementary “Detailed Methods” section.

**Figure 1.**
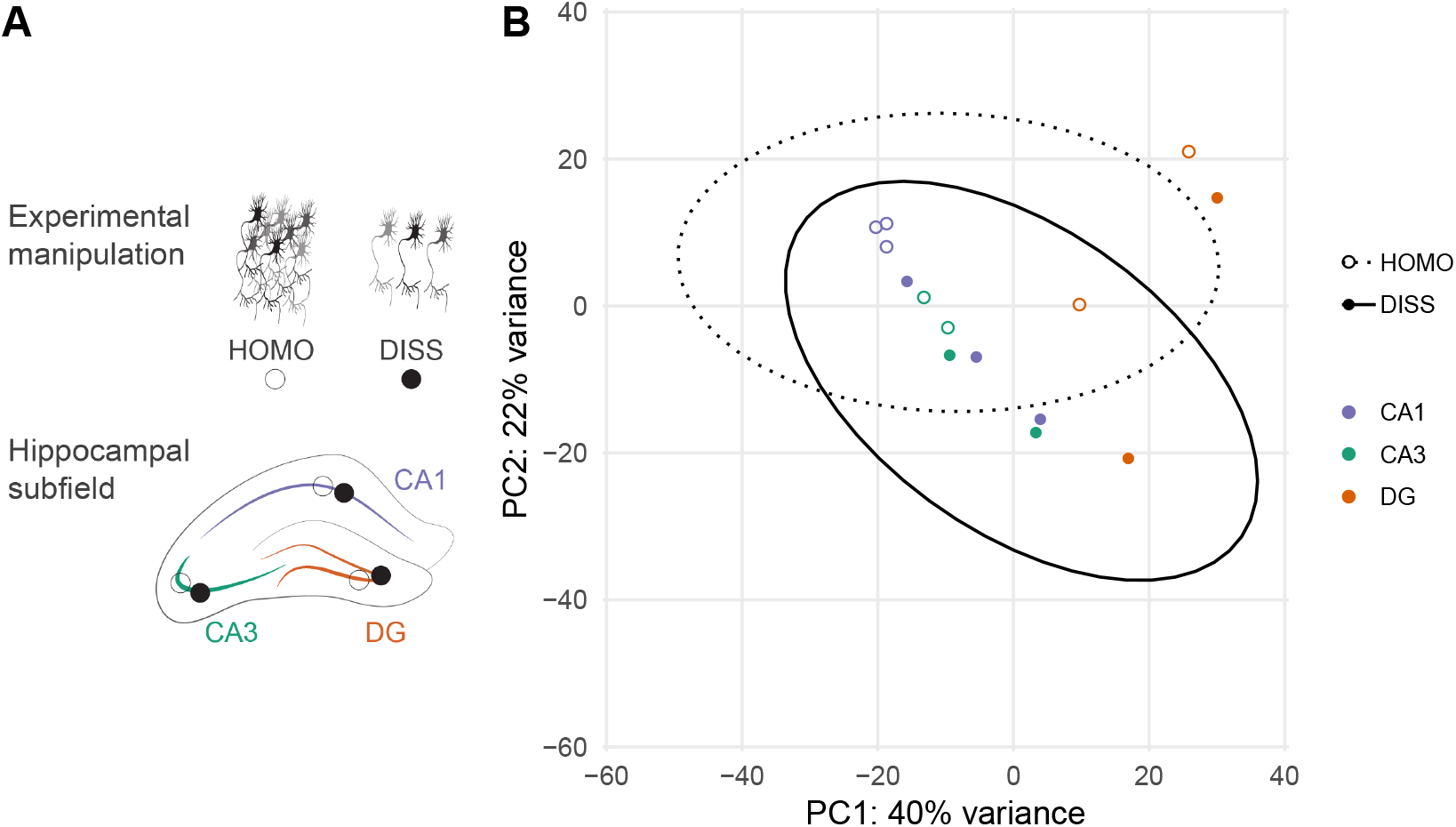
Experimental design and global expression gene expression patterns. **A)** Experimental design. Two tissue samples were taken from three hippocampal subfields (CA1, CA3, and DG) from 300 um brain slices. Two adjacent samples were processed using a homogenization (HOMO) protocol or dissociated (DISS) before processing for tissue level gene expression profiling. **B)** Dissociation does not yield subfield-specific changes in gene expression between homogenized (HOMO, open circles, dotted ellipse) and dissociated tissues (DISS, filled circles, solid ellipse). PC1 accounts for 40% of all gene expression variation and by inspection, separates the DG samples (orange circles) from the CA1 (purple circles) and CA3 samples (green circles). PC2 accounts for 22% of the variation in gene expression and varies significantly with treatment. The ellipses estimate the 95% confidence interval for a multivariate t-distribution for homogenized (dashed line) and dissociated (solid line) samples.

The RNA concentration of samples from homogenized samples (1.45 ± 0.68 ng/*μ*L) was significantly higher than the concentration of samples from dissociated samples (0.48 ± 0.67 ng/*μ*L; F_1_,8 = 7.47, p = 0.026). There was no significant difference in the mean RNA concentration between different subfields (F_2,8_ = 1.15, p = 0.36; or the treatment X subfield interaction F_2,8_ = 0.001, p = 1.0). The number of RNA million reads per sample was not significantly greater in the homogenized (6.30 ± 2.37) compared to the dissociated samples (3.54 ± 2.17; F_1_,8 = 3.81; p = 0.087), nor was there a significant difference in the mean number of reads between different subfields (F_2,8_ = 0.045, p = 0.96) or the interaction between the treatments and subfields (F_2,8_ = 0.38, p = 0.70). On average, 61.2 ± 20.8% of the trimmed reads were pseudoaligned to the mouse transcriptome. Although the sequencing depth was different for each treatment group, this was accounted for by DESeq2, which normalizes counts by sequencing depth to estimate differential gene expression.

The null hypothesis is that treatment effects will not be different between hippocampal subfields. However it is known, that there are subfield expression differences (***Cembrowski et al., 2016a,b, 2018; Hawrylycz et al., 2012; Lein et al., 2004***). DNA microarray followed by in situ hybridization was used to validate region-specific expression patterns of 100 differentially expressed genes (***Lein et al., 2004***). Hierarchical clustering was used to visualize the top 30 differentially expressed genes (p < 0.01) across hippocampal subfields (***Hawrylycz et al., 2012***). RNA-seq experiments on spatially distinct hippocampal subfield samples gave good agreement with immunohistochemical (IHC) data, correctly predicting the enriched populations in 81% of cases (124/153 genes) where coronal IHC images were available (***Cembrowski et al., 2016a***). Because the CA1 region is more vulnerable to anoxia than other hippocampus cell regions (***Pulsinelli et al., 1982***), region-specific differences in the influence of treatment type might also be expected.

We first quantified the effects of treatment and hippocampus subfield on differential gene expression using principal component dimensionality reduction. Samples with similar expression patterns will cluster in the space defined by principal component dimensions. If there are large differences in expression according to treatment, the samples will separate into two non-overlapping clusters. Principal component analysis (PCA) suggests that dissociation does not have a large effect on gene expression because the samples do not form distinct, non-overlapping clusters of homogenized and dissociated samples (***Figure 1***B).

In this analysis the first principal component (PC1) accounts for 40% of the variance and, mostly notably, distinguishes DG samples from the CA1 and CA3 samples. A two-way treatment-by-subfield ANOVA confirmed a significant effect of treatment (F_1_,8 = 5.36, p = 0.049) and subfield (F_2,8_ = 22.48, p = 0.0005) but not the interaction (F_2,8_ = 0.31; p = 0.74). Post hoc Tukey tests confirmed CA1 = CA3 < DG. The second principal component (PC2) accounts for 22% of the variation in gene expression but does not vary significantly with treatment (F_1_,8 = 5.06, p = 0.055), subfield (F_2,8_ = 0.89, p = 0.45), or the interaction (F_2,8_ = 0.062, p = 0.94). None of the higher principal components showed significant variation according to either subfield or treatment. Thus, enzymatic dissociation causes differential gene expression, but the magnitude of the difference is only a fraction of the gene expression differences between hippocampal subfields.

Next, we identified the 344 differentially expressed genes between homogenized and dissociated tissues, accounting for 2.1% of the 16,709 measured genes (***Table 1*** and ??). Most differentially expressed genes showed increased expression (288 genes) rather than decreased expression (56 genes) in response to dissociation (***Figure 2***A). We found that 2.9% of the transcriptome is differentially expressed between CA1 and DG, witha roughly symmetric distribution of differential gene expression (not shown). A heatmap of the top 30 differentially expressed genes illustrates the fold-change differences across samples (***Figure 2***B). Enzymatic dissociation appears to activate gene expression, suggesting the process overall, induces rather than suppresses a cellular response.

**Figure 2.**
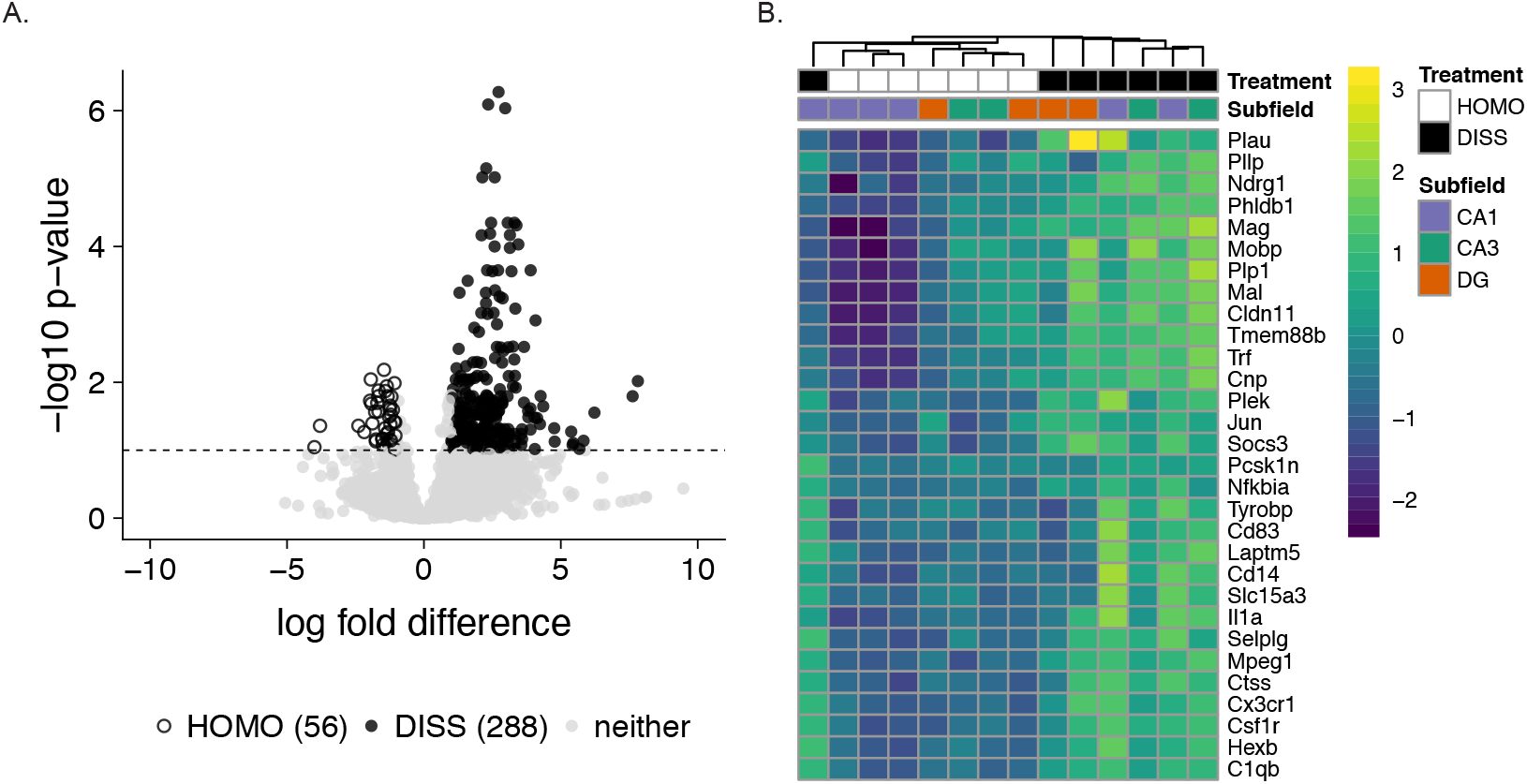
Enzymatic dissociation has a moderate effect on hippocampal gene expression patterns compared to homogenized tissue. **A)** Volcano plot showing gene expression fold-difference and significance between treatment groups. We found that 56 genes are up-regulated in the homogenization control group (open circles) while 288 genes are up-regulated in the dissociated treatment group (filled dark grey circles). Genes below the p-value < 0.1 (or −log p-value < 1) are shown in light grey. **B)** Heatmap showing the top 30 differentially expressed genes between dissociated and homogenized tissue. Square boxes at the top are color coded by sample (white: homogenized, grey: dissociated, purple: CA1, green: CA3, orange: DG. Within the heatmap, log fold difference levels of expression are indicated by the blue-green-yellow gradient with lighter colors indicating increased expression.

**Table 1.**
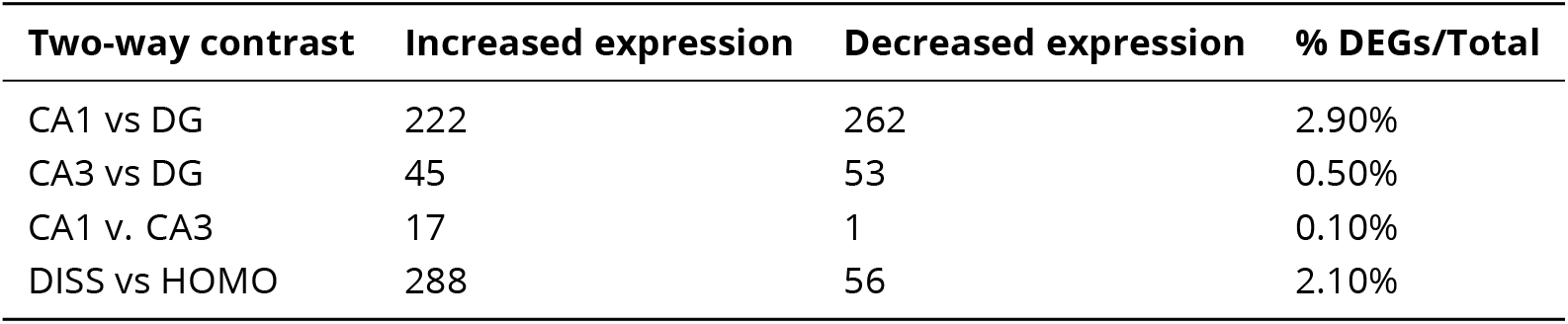
Differentially expressed genes by subfield and treatment. The total number and percent of differentially expressed genes (DEGs) for four two-way contrasts were calculated using DESeq2. Increased expression cutoffs are defined as logfold-change > 0; p < 0.1 while decreased expression is defined as log fold-change < 0; p < 0.1. % DEGs/Total: The sum of up and down regulated genes divided by the total number of genes analyzed (16,709) multiplied by 100%. This table shows that differences between dissociated (DISS) tissue and homogenized (HOMO) tissues are on the same scale as those between the CA1 and DG subfields of the hippocampus.

| Two-way contrast | Increased expression | Decreased expression | % DEGs/Total |
| --- | --- | --- | --- |
| CA1 vs DG | 222 | 262 | 2.90% |
| CA3 vs DG | 45 | 53 | 0.50% |
| CA1 v. CA3 | 17 | 1 | 0.10% |
| DISS vs HOMO | 288 | 56 | 2.10% |

Because the hippocampus is central to learning and memory, we asked whether the expression of genes and pathways known to be involved in learning and memory is affected by dissociation. We first examined expression of 240 genes that have been implicated in long-term potentiation (LTP) (***Sanes and Lichtman, 1999***) ?? and found that the expression of only nine of these genes was altered by enzymatic dissociation treatment. The expression of *CACNA1E, GABRB1, GRIN2A* was downregulated in response to dissociation treatment (meaning that their activity could be underestimated in an experiment using enzymatic treatment to dissociate tissue) while ***IL1B, ITGA5, ITGAM, ITGB4, ITGB5, and MAPK3*** were upregulated in response to dissociation. CACNA1E is a subunit of L-type calcium channels, which are necessary for LTP induction of mossy fiber input to CA3 pyramidal neurons (***Kapur et al., 1998***). *GABRB1* encodes the Gamma-Aminobutyric Acid (GABA) A Receptor Beta subunit, and *GRIN2A* encodes the Glutamate Ionotropic Receptor NMDA Type 2A subunit. Because GABA receptors and NMDA receptors mediate inhibitory and excitatory neurotransmission in hippocampus, respectively, enzymatic dissociation could itself alter accurate estimation of the roles of these receptors. IL1B encodes interleukin-1beta, a cytokine that plays a key role in the immune response to infection and injury but is also critical for maintaining LTP in heathy brains (***Schneider et al., 1998***). The integrin class of cell adhesion molecules plays an important role in synaptic plasticity, particularly in stabilization and consolidation of LTP (***Bahr et al., 1997; McGeachie et al.,2011***). Overall, our analysis demonstrates that the expression of only a few cannonical LTP-related genes is affected by the tissue prepraration method.

More recently, RNA sequencing was used in combination with ribosomal profiling to quantify the translational status and transcript levels in the mouse hippocampus after contextual fear conditioning (***Cho et al., 2015***). The analysis revealed that memory formation was regulated by learning-induced suppression of ribosomal protein-coding genes and suppression of a subset of genes via inhibition of estrogen receptor 1 signaling in the hippocampus. We cross-referenced learning-induced differential gene expression from (***Cho et al., 2015***), to identify genes that are altered by both fear-conditioning and enzymatic dissociation. We found that *BTG2, FOSB, FN1, IER2*, and *JUNB* were all upregulated in response to enzymatic dissociation and fear-conditioning while *Enpp2* was upregulated in response to dissociation but down-regulated in fear-conditioning via estrogen receptor 1 inhibition. *BTG2* is required for proliferation and differentiation of neurons during adult hippocampal neurogenesis and may be involved in the formation of contextual memories ***Farioli-Vecchioli et al. (2009)***. FOSB and JUNB are dimers that form the transcription factor complex AP-1 that is often used as a marker for neural activity (***Alberini, 2009***). *IER2* is also a transcription factor that, along with FOS and JUN, as well as *FN1*, which encodes the adhesion molecule Fibronectin, was not included in the (***Sanes and Lichtman, 1999***) list as important for LTP but was differentially expressed following fear-conditioning in (***Cho et al., 2015***). These comparisons show that tissue preparation methods can alter expression in a small subset of genes that may be important for LTP.

This study was motivated by the possibility of single cell sequencing, although we did not conduct single-neuron sequencing in this study. A single-cell study would not have made it possible to test our hypothesis of how the process of cellular dissociation affects gene expression relative to tissue homogenization, because the RNA from single cells can’t be recovered after tissue homogenization. To compare single cell transcriptomes that are obtained without dissociation, we could have used mechanical dissociation for example by laser microdissection and capture or by microaspiration but this was not deemed practical because these are substantially more difficult, expensive, and low-throughput procedures compared to enzymatic dissociation of cells. Given the present findings that enzymatic dissociation may itself induce gene expression, it may be useful to first prepare tissues with transcription and translation blockers like puromycin and actinomycin to arrest gene expression activity before cellular dissociation (***Flexner et al., 1963; Solntseva and Nikitin, 2012***), but potential additional effects of these treatments will also need to be investigated and controlled using appropriate experimental designs.

We set out to identify the extent to which the process of chemical cellular dissociation, affects neural gene expression profiles because the process necessarily precedes high-throughput single cell analysis of complex tissues. One possible confounding factor is that the process of dissociation could kill some cell classes in the hippocampus, either indiscriminately or preferentially, which could explain the lower RNA content after the dissociation treatment. Accordingly, we examined whether well-described marker genes for astrocytes, oligodendrocytes, microglia, and neurons were over- or under-expressed in the dissociated samples compared to the homogenized samples (***Cahoy et al., 2008***). None of the marker genes for astrocytes or neurons was differentially expressed, but 1 of 3 and 7 of 10 markers for microglia and oligodendrocytes, respectively, were over-expressed in the dissociated samples (??). This overexpression could arise if these cells were more resilient during the dissociation. Because neural makers were not over-expressed in the homogenized tissue, it is unlikely that dissociation preferentially kills neurons.

In summary, we found that gene expression in hippocampal subfields is changed by tissue preparation procedures (cellular dissociation versus homogenization) and cross-referenced the differentially expressed genes with genes and pathways known to be involved in hippocampal LTP, learning and memory. While it is encouraging that the activity of only a small number of genes and pathways involved in LTP, learning and memory appears affected by dissociation, it is also important to effectively use experimental design to control for technical artifacts. The present findings provide insight into how cellular manipulations influence gene expression, which is important because it is increasingly necessary to dissociate cells in tissue samples for single cell or single cell-type studies.

## Acknowledgments

We thank members of the Hofmann and Fenton Labs, Boris Zemelman, Laura Colgin, and Misha Matz for helpful discussions. We thank Dennis Wylie for insightful comments on earlier versions of this manuscript. We thank the GSAF for library preparation and sequencing. The bioinformatic workflow was inspired heavily by Center for Computational Biology’s Bioinformatics Curriculum and Software Carpentry Curriculum on the Unix Shell, Git for Version Control, and R for Reproducible Research. This work is supported by a Society for Integrative Biology (SICB) Grant in Aid of Research (GIAR) grant and a UT Austin Graduate School Continuing Fellowship to RMH; a generous gift from Michael Vasinkevich to AAF; NIH-NS091830 to JMA, IOS-1501704 to HAH; NIMH-5R25MH059472-18. The authors declare no competing interests.

## Detailed methods

All animal care and use comply with the Public Health Service Policy on Humane Care and Use of Laboratory Animals and were approved by the New York University Animal Welfare Committee. A1-year-old female C57BL/6J mouse was taken from its cage, anesthetized with 2% (vol/vol) isoflurane for 2 minutes and decapitated. Transverse 300 μm brain slices were cut using a vibratome (model VT1000 S, Leica Biosystems, Buffalo Grove, IL) and incubated at 36°C for 30 min and then at room temperature for 90 min in oxygenated artificial cerebrospinal fluid (aCSF in mM: 125 NaCl, 2.5 KCl, 1 MgSO4, 2 CaCl2, 25 NaHCO3, 1.25 NaH2PO4 and 25 Glucose) as in Pavlowsky and Alarcon, 2012. Tissue adjacent samples were collected from CA1, CA3, and DG, respectively in the dorsal hippocampus by punch (0.25 mm, P/N: 57391; Electron Microscopy Sciences, Hatfield, PA) (Fig 1A).

The homogenized (HOMO) samples were processed using the manufacturer instructors for the Maxwell 16 LEV RNA Isolation Kit (Promega, Madison, WI). The dissociated (DISS) samples were incubated for 75 minutes in aCSF containing 1 mg/ml pronase at room temperature, then vortexed and centrifuged. The incubation was terminated by replacing aCSF containing pronase with aCSF. The sample was then vortexed, centrifuged, and gently triturated by 200-*μ*L pipette tip twenty times in aCSF containing 1% FBS. The sample was centrifuged and used as input for RNA isolation using the Maxwell 16 LEV RNA Isolation Kit (Promega, Madison, WI).

RNA libraries were prepared by the Genomic Sequencing and Analysis Facility at the University of Texas at Austin using the Illumina HiSeq platform. Raw reads were processed and analyzed on the Stampede Cluster at the Texas Advanced Computing Facility (TACC). Samples yielded an average of 4.9 +/− 2.6 million reads. Read quality was checked using the program FASTQC. Low quality reads and adapter sequences were removed using the program Cutadapt (***Martin, 2011***). We used Kallisto for read pseudoalignment to the Gencode M11 mouse transcriptome and for transcript counting (***Bray et al., 2016; Mudge and Harrow, 2015***). On average, 61.2% +/− 20.8% of the trimmed reads were pseudoaligned to the mouse transcriptome. Two-way ANOVAs were used to test for significant differences (p-value < 0.5) in RNA concentration and read counts for treatment and subfield.

Kallisto transcript counts were imported into R (***R Development Core Team, 2013***) and aggregated to yield gene counts using the ‘gene’ identifier from the Gencode reference transcriptome. We used DESeq2 for gene expression normalization and quantification of gene level counts (***Love et al., 2014***). We used a threshold of a false discovery corrected (FDR) p-value < 0.1. Statistics on the principal component analysis (PCA) were conducted in R. The hierarchical clustering analysis was conducted and visualized using the R package pheatmap (***Kolde, 2015***) with the RColorBrewer R packages for color modifications (***Neuwirth, 2014***). PCA was conducted in R using the DESeq2 and genefilter R packages (***Gentleman R et al., 2017; Love et al., 2014***) and visualized using the ggplot2 and cowplot R packages (***Wilke, 2016; Wickham, 2009***). Two-way ANOVAs were used to test whether or not a significant amount of variance in PC1 and PC2 is explained by treatment, subfield, or their interaction.

The raw sequence data and intermediate data files are archived in NCBI’s Gene Expression Omnibus Database (accession numbers GSE99765). The data and code are available on GitHub (https://github.com/raynamharris/DissociationTest), with an archived version at the time of publication available at Zenodo (Harris et al., 2017). A Jupyter notebook containing a cloud-based, open-access analysis of GEO dataset GSE99765 (https://www.ncbi.nlm.nih.gov/gds/?term=GSE99765) created using BioJupies (***Torre et al., 2018***) is available at http://amp.pharm.mssm.edu/biojupies/notebook/zySloEXuZ.

## Supplementary Materials

**Table 2.**
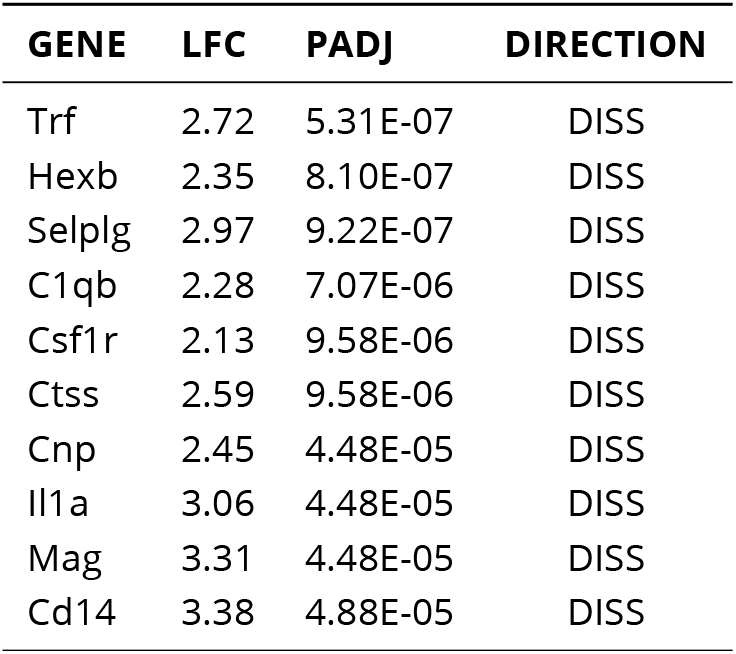
Expression level and fold change of significant genes (p < 0.1) between dissociated tissue and homogenized tissue. This table shows the log fold change (LFC), adjusted p-value (PADJ), and direction of increased expression (DISS, HOMO, or neither) for each gene analyzed. The full table is available at https://github.com/raynamharris/DissociationTest/blob/master/results/dissociationDEGs.csv.

**Table 3.**
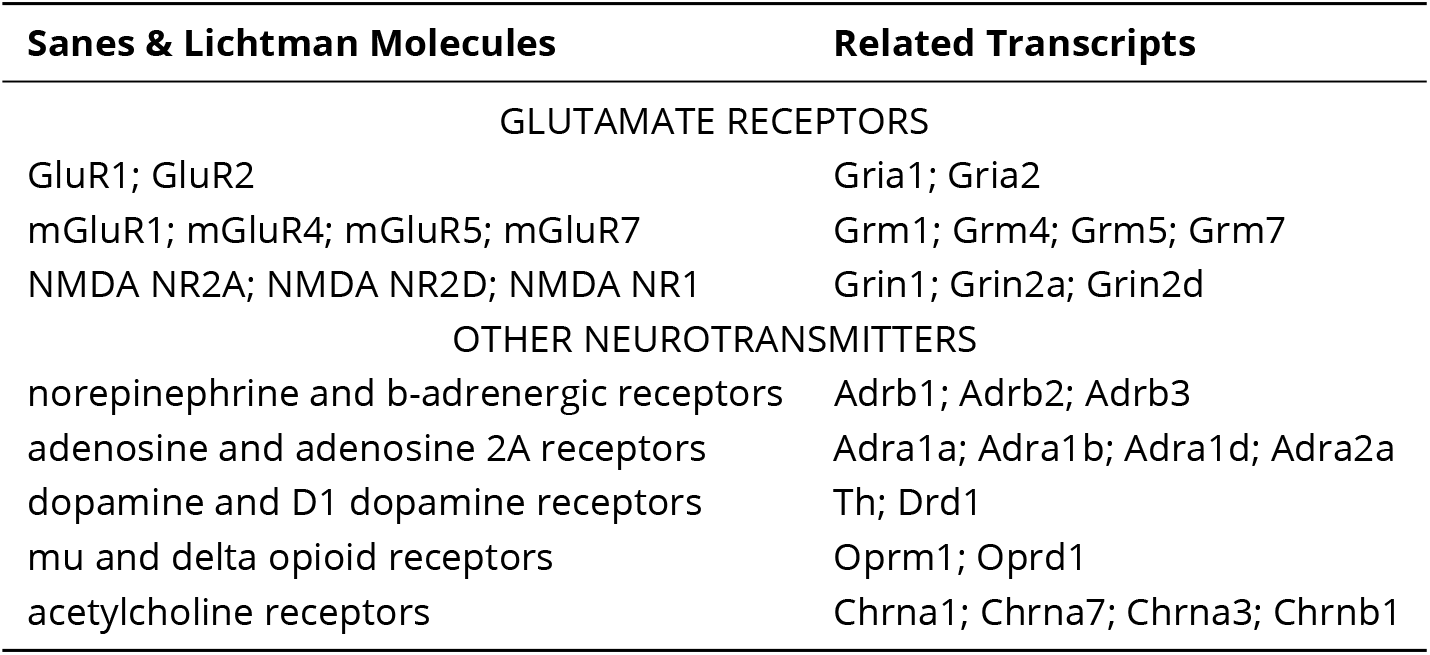
Molecules implicated in hippocampal LTP from Sanes and Lichtman 1999. This table list the molecules review by Sanes and Lichtman in their 1999 review article and the related transcripts that were investigated in this study. *This is a preview. The full table is available at* https://github.com/raynamharris/DissociationTest/blob/master/data/SanesLichtman.csv

**Table 4.**
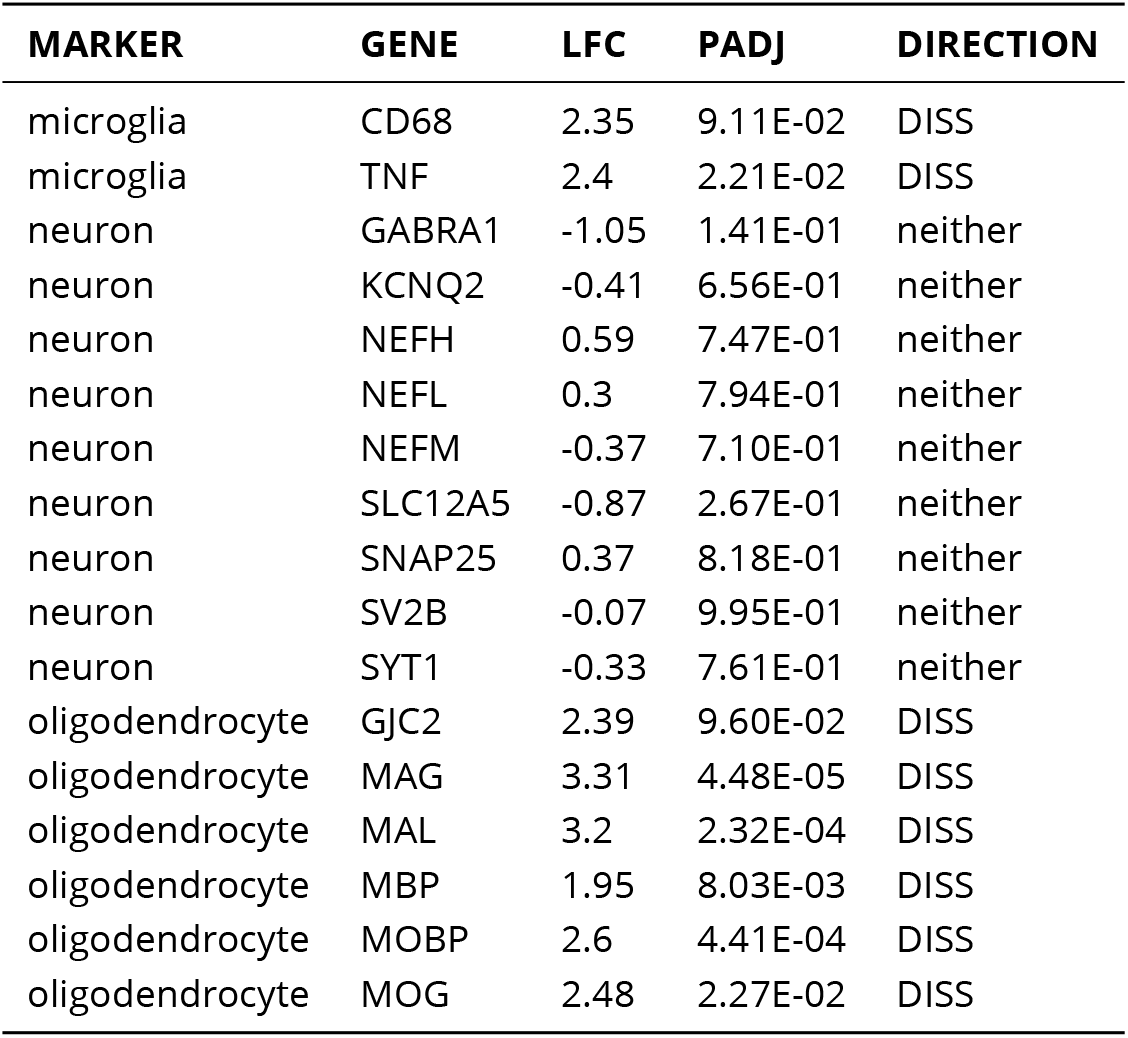
Marker genes for astrocytes, oligodendrocytes, microglia, and neurons. This table, adapted from Cahoy et al., 2008, lists the genes we investigated to estimate the relative abundance of cell types in the examined tissue. LFC: Log fold change; PADJ: adjusted p-value; DIRECTION: upregulated in dissociated (DISS) or not up-regulated in either dissociated or homogenized (neither)

