## Supplementary material for "Hippocampal transcriptomic responses to cellular dissociation": Supp. Table 1

| gene | lfc | padj | direction |
| --- | --- | --- | --- |
| *Trf* | 2.72 | 5.31E-07 | DISS |
| *Hexb* | 2.35 | 8.10E-07 | DISS |
| *Selplg* | 2.97 | 9.22E-07 | DISS |
| *C1qb* | 2.28 | 7.07E-06 | DISS |
| *Csf1r* | 2.13 | 9.58E-06 | DISS |
| *Ctss* | 2.59 | 9.58E-06 | DISS |
| *Cnp* | 2.45 | 4.48E-05 | DISS |
| *Il1a* | 3.06 | 4.48E-05 | DISS |
| *Mag* | 3.31 | 4.48E-05 | DISS |
| *Cd14* | 3.38 | 4.88E-05 | DISS |
| *Mpeg1* | 2.42 | 6.47E-05 | DISS |
| *Tmem88b* | 3.14 | 6.78E-05 | DISS |
| *Nfkbia* | 2.1 | 6.84E-05 | DISS |
| *Slc15a3* | 3.45 | 9.32E-05 | DISS |
| *Cd83* | 2.58 | 9.99E-05 | DISS |
| *Cldn11* | 3.14 | 1.05E-04 | DISS |
| *Laptm5* | 2.31 | 2.24E-04 | DISS |
| *Plau* | 3.9 | 2.24E-04 | DISS |
| *Plp1* | 2.71 | 2.24E-04 | DISS |
| *Mal* | 3.2 | 2.32E-04 | DISS |
| *Plek* | 2.5 | 2.32E-04 | DISS |
| *Jun* | 1.6 | 3.20E-04 | DISS |
| *Mobp* | 2.6 | 4.41E-04 | DISS |
| *Pcsk1n* | 1.3 | 4.80E-04 | DISS |
| *Socs3* | 2.26 | 4.80E-04 | DISS |
| *Pllp* | 2.76 | 5.46E-04 | DISS |
| *Ndrg1* | 2.87 | 5.80E-04 | DISS |
| *Phldb1* | 2.27 | 6.85E-04 | DISS |
| *Tyrobp* | 3.34 | 8.22E-04 | DISS |
| *Cx3cr1* | 2.1 | 9.50E-04 | DISS |
| *Tmem119* | 2.54 | 9.50E-04 | DISS |
| *C1qa* | 2.32 | 9.85E-04 | DISS |
| *Slc2a5* | 4.08 | 1.23E-03 | DISS |
| *Fcrls* | 2.67 | 1.40E-03 | DISS |
| *Lgmn* | 1.84 | 1.57E-03 | DISS |
| *Clic4* | 2.01 | 1.81E-03 | DISS |
| *Sema5a* | 3.25 | 2.98E-03 | DISS |
| *C5ar1* | 3.66 | 3.01E-03 | DISS |
| *Cmtm5* | 2.66 | 3.01E-03 | DISS |
| *Ctcfl* | 3.08 | 3.05E-03 | DISS |
| *Cryab* | 2.81 | 3.07E-03 | DISS |
| *Rpl26* | 1.27 | 3.22E-03 | DISS |
| *Gab1* | 2.9 | 3.44E-03 | DISS |
| *C1qc* | 2.6 | 4.40E-03 | DISS |
| *Fa2h* | 3.3 | 4.61E-03 | DISS |
| *Clec2l* | 1.94 | 5.01E-03 | DISS |
| *Spry2* | 1.79 | 5.08E-03 | DISS |
| *Mcam* | 2.86 | 5.12E-03 | DISS |
| *Olfml3* | 2.09 | 5.18E-03 | DISS |
| *Sepp1* | 1.54 | 5.78E-03 | DISS |
| *Ptma* | 1.19 | 6.21E-03 | DISS |
| *Csrnp3* | -1.46 | 6.55E-03 | HOMO |
| *Dusp1* | 1.35 | 8.03E-03 | DISS |
| *Lag3* | 3.32 | 8.03E-03 | DISS |
| *Mbp* | 1.95 | 8.03E-03 | DISS |
| *Atf3* | 2.17 | 8.09E-03 | DISS |
| *C1ql1* | 3.1 | 8.09E-03 | DISS |
| *Adgrg1* | 1.86 | 8.62E-03 | DISS |
| *Csrp1* | 1.7 | 9.07E-03 | DISS |
| *Ltbp3* | 2.29 | 9.07E-03 | DISS |
| *Mcl1* | 1.41 | 9.07E-03 | DISS |
| *Rpl29* | 1.16 | 9.07E-03 | DISS |
| *Rpl36a* | 1.57 | 9.07E-03 | DISS |
| *Stxbp5l* | -1.94 | 9.07E-03 | HOMO |
| *Rps27* | 1.29 | 9.59E-03 | DISS |
| *Ighv14-2* | 7.81 | 9.60E-03 | DISS |
| *Mapk3* | 1.61 | 9.80E-03 | DISS |
| *Gabrb1* | -1.07 | 1.03E-02 | HOMO |
| *Rgs7bp* | -1.36 | 1.14E-02 | HOMO |
| *Trem2* | 3.25 | 1.14E-02 | DISS |
| *Syngr2* | 3.35 | 1.20E-02 | DISS |
| *Irf8* | 2.77 | 1.26E-02 | DISS |
| *Jund* | 1.06 | 1.26E-02 | DISS |
| *Plekhb1* | 1.49 | 1.26E-02 | DISS |
| *Grin2b* | -1.66 | 1.32E-02 | HOMO |
| *Sorl1* | -1.41 | 1.35E-02 | HOMO |
| *Cst3* | 1.53 | 1.45E-02 | DISS |
| *Rpl28* | 1.17 | 1.55E-02 | DISS |
| *Unc93b1* | 2.28 | 1.55E-02 | DISS |
| *D6Wsu163e* | 2.32 | 1.59E-02 | DISS |
| *Sstr1* | 4.26 | 1.59E-02 | DISS |
| *Birc6* | -1.32 | 1.60E-02 | HOMO |
| *Cdkl5* | -1.67 | 1.60E-02 | HOMO |
| *Gm10401* | 7.62 | 1.60E-02 | DISS |
| *Itga5* | 3.05 | 1.60E-02 | DISS |
| *Ly86* | 2.81 | 1.60E-02 | DISS |
| *Olig1* | 1.67 | 1.60E-02 | DISS |
| *Zfp36* | 1.79 | 1.60E-02 | DISS |
| *Ralgapa1* | -1.19 | 1.63E-02 | HOMO |
| *Rps8* | 1.04 | 1.70E-02 | DISS |
| *Cd9* | 2.36 | 1.72E-02 | DISS |
| *Rph3a* | 2.26 | 1.72E-02 | DISS |
| *Slc2a1* | 1.7 | 1.77E-02 | DISS |
| *Bin2* | 2.49 | 1.83E-02 | DISS |
| *Gpr84* | 2.86 | 1.87E-02 | DISS |
| *H2-K1* | 2.09 | 1.87E-02 | DISS |
| *Rc3h2* | -1.97 | 1.87E-02 | HOMO |
| *Slco2b1* | 2.17 | 1.87E-02 | DISS |
| *Tgfbr2* | 2.44 | 1.87E-02 | DISS |
| *Cdh9* | 3.65 | 1.99E-02 | DISS |
| *Ctsd* | 1.26 | 1.99E-02 | DISS |
| *Ptgds* | 2.53 | 1.99E-02 | DISS |
| *Epha6* | -1.7 | 2.01E-02 | HOMO |
| *Mt1* | 1.42 | 2.01E-02 | DISS |
| *Itgb5* | 1.98 | 2.01E-02 | DISS |
| *Inpp5d* | 2.84 | 2.04E-02 | DISS |
| *Icam1* | 2.87 | 2.06E-02 | DISS |
| *Kcnma1* | -1.92 | 2.06E-02 | HOMO |
| *Myrf* | 2.21 | 2.21E-02 | DISS |
| *Nos1ap* | -1.23 | 2.21E-02 | HOMO |
| *Sparc* | 1.79 | 2.21E-02 | DISS |
| *Tnf* | 2.4 | 2.21E-02 | DISS |
| *Siglech* | 2.11 | 2.21E-02 | DISS |
| *Serpinb1a* | 4.36 | 2.26E-02 | DISS |
| *mt-Co3* | 1.4 | 2.26E-02 | DISS |
| *Mog* | 2.48 | 2.27E-02 | DISS |
| *Stat3* | 1.56 | 2.41E-02 | DISS |
| *Tlr7* | 3.89 | 2.42E-02 | DISS |
| *Plppr4* | -1.25 | 2.49E-02 | HOMO |
| *Celsr2* | -1.13 | 2.54E-02 | HOMO |
| *Lgals9* | 2.74 | 2.62E-02 | DISS |
| *Tln1* | 1.34 | 2.62E-02 | DISS |
| *Cyba* | 3.8 | 2.63E-02 | DISS |
| *Ermn* | 2.53 | 2.69E-02 | DISS |
| *Fn1* | 1.77 | 2.69E-02 | DISS |
| *Grin2a* | -1.66 | 2.69E-02 | HOMO |
| *mt-Co1* | 1.24 | 2.69E-02 | DISS |
| *Sipa1* | 2.71 | 2.69E-02 | DISS |
| *Lrrc7* | -1.78 | 2.73E-02 | HOMO |
| *Vasp* | 2.31 | 2.73E-02 | DISS |
| *Shroom3* | 6.23 | 2.80E-02 | DISS |
| *mt-Cytb* | 1.27 | 2.86E-02 | DISS |
| *Qdpr* | 1.55 | 2.86E-02 | DISS |
| *Slc12a2* | 2.07 | 2.86E-02 | DISS |
| *Tmem170* | 2.53 | 2.93E-02 | DISS |
| *Apod* | 2.05 | 3.00E-02 | DISS |
| *Cadm2* | -1.21 | 3.00E-02 | HOMO |
| *Ccl4* | 1.94 | 3.00E-02 | DISS |
| *Erdr1* | 1.33 | 3.00E-02 | DISS |
| *Il1b* | 2.41 | 3.00E-02 | DISS |
| *Plekhg3* | 2.65 | 3.00E-02 | DISS |
| *Sep4* | 1.84 | 3.00E-02 | DISS |
| *Cd82* | 2.18 | 3.20E-02 | DISS |
| *Cspg5* | 1.28 | 3.21E-02 | DISS |
| *Magi2* | -1.27 | 3.21E-02 | HOMO |
| *Ucp2* | 2.49 | 3.21E-02 | DISS |
| *Smoc2* | 3.81 | 3.27E-02 | DISS |
| *mt-Nd3* | 1.7 | 3.28E-02 | DISS |
| *Fosb* | 1.59 | 3.32E-02 | DISS |
| *Mfng* | 4.05 | 3.32E-02 | DISS |
| *Pcdhga8* | 1.42 | 3.32E-02 | DISS |
| *Emp2* | 2.35 | 3.33E-02 | DISS |
| *Cd37* | 4.14 | 3.36E-02 | DISS |
| *Scrg1* | 2.63 | 3.36E-02 | DISS |
| *Prr18* | 2.29 | 3.57E-02 | DISS |
| *mt-Nd1* | 1.21 | 3.60E-02 | DISS |
| *Rasgrp1* | -1.08 | 3.78E-02 | HOMO |
| *Rock2* | -1.05 | 3.88E-02 | HOMO |
| *Asap1* | -1.88 | 4.02E-02 | HOMO |
| *Cd274* | 4.24 | 4.12E-02 | DISS |
| *Rpl32* | 1.33 | 4.12E-02 | DISS |
| *Synpr* | 2.31 | 4.12E-02 | DISS |
| *Cacna2d1* | -1.22 | 4.17E-02 | HOMO |
| *Pold1* | 2.23 | 4.17E-02 | DISS |
| *Btg2* | 1.39 | 4.21E-02 | DISS |
| *Gm42878* | -3.8 | 4.37E-02 | HOMO |
| *Kcnj6* | -2.39 | 4.37E-02 | HOMO |
| *Crtc2* | 1.78 | 4.37E-02 | DISS |
| *Msn* | 1.75 | 4.55E-02 | DISS |
| *mt-Nd4* | 1.15 | 4.55E-02 | DISS |
| *Rtn2* | 1.47 | 4.55E-02 | DISS |
| *Myc* | 2.35 | 4.56E-02 | DISS |
| *Bcas1* | 1.7 | 4.70E-02 | DISS |
| *Adcy9* | -1.42 | 4.74E-02 | HOMO |
| *Gatm* | 2.12 | 4.74E-02 | DISS |
| *Gjb1* | 4.76 | 4.74E-02 | DISS |
| *Pros1* | 2.29 | 4.74E-02 | DISS |
| *Psat1* | 1.31 | 4.74E-02 | DISS |
| *Rab3il1* | 2.76 | 4.78E-02 | DISS |
| *Arl4c* | 1.79 | 4.89E-02 | DISS |
| *Dhrs3* | 2.95 | 4.89E-02 | DISS |
| *Enpp2* | 1.77 | 4.89E-02 | DISS |
| *Gpr34* | 2.08 | 4.89E-02 | DISS |
| *Ntng1* | 3.08 | 4.89E-02 | DISS |
| *P2ry12* | 1.74 | 4.89E-02 | DISS |
| *Tlr2* | 2.65 | 4.89E-02 | DISS |
| *Cyp27a1* | 3.58 | 4.92E-02 | DISS |
| *Gna12* | 1.3 | 4.92E-02 | DISS |
| *Asap3* | 3.89 | 4.96E-02 | DISS |
| *Itgam* | 1.75 | 4.96E-02 | DISS |
| *Lcp1* | 2.73 | 4.96E-02 | DISS |
| *Frmd8* | 2.79 | 5.03E-02 | DISS |
| *Ncf1* | 1.8 | 5.07E-02 | DISS |
| *Dnajb2* | 1.52 | 5.14E-02 | DISS |
| *mt-Atp6* | 1.25 | 5.15E-02 | DISS |
| *mt-Co2* | 1.36 | 5.15E-02 | DISS |
| *Wasf2* | 1.67 | 5.20E-02 | DISS |
| *Ccdc3* | 5.38 | 5.32E-02 | DISS |
| *Fat3* | -2.18 | 5.40E-02 | HOMO |
| *Smox* | 1.58 | 5.40E-02 | DISS |
| *Taok1* | -1.31 | 5.40E-02 | HOMO |
| *Tmem63a* | 2.46 | 5.40E-02 | DISS |
| *Vsir* | 2.06 | 5.40E-02 | DISS |
| *Gpr17* | 2.6 | 5.46E-02 | DISS |
| *Adgrl3* | -1.35 | 5.70E-02 | HOMO |
| *Cmtm6* | 1.7 | 5.75E-02 | DISS |
| *Dock8* | 3.07 | 5.75E-02 | DISS |
| *Gp1bb* | 3.1 | 5.75E-02 | DISS |
| *Ier2* | 1.33 | 5.75E-02 | DISS |
| *Il16* | 3.32 | 5.75E-02 | DISS |
| *Rftn1* | 3.55 | 5.75E-02 | DISS |
| *Sox10* | 2.16 | 5.75E-02 | DISS |
| *Junb* | 1.03 | 5.81E-02 | DISS |
| *D630003M21Rik* | 3.12 | 5.89E-02 | DISS |
| *Kcnk6* | 2.84 | 6.10E-02 | DISS |
| *Slc4a8* | -1.04 | 6.10E-02 | HOMO |
| *Rela* | 1.52 | 6.18E-02 | DISS |
| *mt-Nd4l* | 1.06 | 6.21E-02 | DISS |
| *Itgb4* | 2.93 | 6.30E-02 | DISS |
| *Zfp521* | 2.72 | 6.30E-02 | DISS |
| *Ilkap* | 1.61 | 6.31E-02 | DISS |
| *Nckap1l* | 2.11 | 6.31E-02 | DISS |
| *Ccrl2* | 3.01 | 6.48E-02 | DISS |
| *Gjc3* | 1.85 | 6.48E-02 | DISS |
| *Csrnp1* | 1.52 | 6.62E-02 | DISS |
| *Klf4* | 1.74 | 6.62E-02 | DISS |
| *Tbc1d16* | 1.48 | 6.65E-02 | DISS |
| *Wscd1* | 1.5 | 6.69E-02 | DISS |
| *Atxn1* | -1.52 | 6.74E-02 | HOMO |
| *Arfgef3* | -1.52 | 6.84E-02 | HOMO |
| *Arhgap32* | -1.28 | 6.85E-02 | HOMO |
| *Atf5* | 1.58 | 6.85E-02 | DISS |
| *Dusp10* | 2.49 | 6.85E-02 | DISS |
| *Prdm5* | 3.59 | 6.85E-02 | DISS |
| *Arpc1b* | 2.05 | 6.89E-02 | DISS |
| *mt-Nd2* | 1.19 | 6.95E-02 | DISS |
| *Pde4d* | -1.75 | 6.95E-02 | HOMO |
| *Grap* | 2.96 | 7.08E-02 | DISS |
| *Rpl18* | 1.04 | 7.08E-02 | DISS |
| *Suv420h2* | 2.25 | 7.08E-02 | DISS |
| *Taf1* | -1.21 | 7.15E-02 | HOMO |
| *Il6ra* | 2.21 | 7.21E-02 | DISS |
| *Npr1* | 3.34 | 7.24E-02 | DISS |
| *Clock* | -1.24 | 7.25E-02 | HOMO |
| *Smtnl2* | 5.83 | 7.25E-02 | DISS |
| *mt-Atp8* | 1.11 | 7.37E-02 | DISS |
| *Piezo1* | 3.06 | 7.37E-02 | DISS |
| *Rpl23a* | 1 | 7.37E-02 | DISS |
| *Rpl31* | 1.09 | 7.37E-02 | DISS |
| *Pim1* | 2.41 | 7.37E-02 | DISS |
| *Atp2b1* | -1.21 | 7.40E-02 | HOMO |
| *Cd63* | 1.58 | 7.40E-02 | DISS |
| *Csf1* | 1.88 | 7.40E-02 | DISS |
| *Prr36* | -1.76 | 7.40E-02 | HOMO |
| *Rplp1* | 1.13 | 7.40E-02 | DISS |
| *Myo1f* | 4.78 | 7.47E-02 | DISS |
| *Insig1* | 1.61 | 7.50E-02 | DISS |
| *Itpk1* | 1.53 | 7.58E-02 | DISS |
| *Lhfpl2* | 2.15 | 7.69E-02 | DISS |
| *Chadl* | 1.82 | 7.69E-02 | DISS |
| *Nlrp3* | 2.2 | 7.69E-02 | DISS |
| *Sh3gl1* | 1.56 | 7.69E-02 | DISS |
| *Cidea* | 5.46 | 7.85E-02 | DISS |
| *Rack1* | 1.03 | 7.85E-02 | DISS |
| *Sema4g* | 2.3 | 7.95E-02 | DISS |
| *Rpl23* | 1.23 | 8.02E-02 | DISS |
| *Adam17* | 2.27 | 8.05E-02 | DISS |
| *Cacna1e* | -1.28 | 8.05E-02 | HOMO |
| *Cpm* | 2.85 | 8.15E-02 | DISS |
| *Cox4i1* | 1.04 | 8.27E-02 | DISS |
| *Nectin1* | 1.84 | 8.27E-02 | DISS |
| *Ostf1* | 2.36 | 8.27E-02 | DISS |
| *Gnaq* | -1.18 | 8.28E-02 | HOMO |
| *Maml2* | 2.19 | 8.28E-02 | DISS |
| *Syt14* | -1.5 | 8.35E-02 | HOMO |
| *Ap1s1* | 1.07 | 8.38E-02 | DISS |
| *Fam124a* | 2.36 | 8.38E-02 | DISS |
| *Hr* | 2.45 | 8.38E-02 | DISS |
| *Nfatc1* | 3.56 | 8.38E-02 | DISS |
| *Rps6kb2* | 1.65 | 8.38E-02 | DISS |
| *Stk32b* | 5.43 | 8.39E-02 | DISS |
| *Unc5b* | 1.96 | 8.52E-02 | DISS |
| *Cdkn1a* | 2.51 | 8.54E-02 | DISS |
| *Fam102b* | 1.53 | 8.75E-02 | DISS |
| *Garem* | -1.39 | 8.88E-02 | HOMO |
| *Pde4c* | -4 | 9.01E-02 | HOMO |
| *Sh3d19* | 2.43 | 9.07E-02 | DISS |
| *Tspan2* | 1.93 | 9.07E-02 | DISS |
| *Tnfaip6* | 2.76 | 9.09E-02 | DISS |
| *Cd68* | 2.35 | 9.11E-02 | DISS |
| *Mmp15* | 1.94 | 9.19E-02 | DISS |
| *Lamb2* | 2.7 | 9.21E-02 | DISS |
| *Itpkb* | 1.53 | 9.25E-02 | DISS |
| *Il10ra* | 2.49 | 9.29E-02 | DISS |
| *Rps27a* | 1.13 | 9.33E-02 | DISS |
| *Rpl35a* | 1.47 | 9.45E-02 | DISS |
| *Tbx1* | 5.68 | 9.56E-02 | DISS |
| *Srgn* | 2.41 | 9.59E-02 | DISS |
| *Apoe* | 1.25 | 9.60E-02 | DISS |
| *Atp2c1* | -1.13 | 9.60E-02 | HOMO |
| *Bche* | 3.35 | 9.60E-02 | DISS |
| *Cacnb2* | -1.16 | 9.60E-02 | HOMO |
| *Ccl9* | 3.31 | 9.60E-02 | DISS |
| *Cd81* | 1.03 | 9.60E-02 | DISS |
| *Cfh* | 2.03 | 9.60E-02 | DISS |
| *Ctsc* | 2.4 | 9.60E-02 | DISS |
| *Dusp26* | 2.02 | 9.60E-02 | DISS |
| *Erbin* | 1.72 | 9.60E-02 | DISS |
| *Gjc2* | 2.39 | 9.60E-02 | DISS |
| *Lrtm2* | 1.8 | 9.60E-02 | DISS |
| *Plin4* | 4.05 | 9.60E-02 | DISS |
| *Rgs10* | 2.18 | 9.60E-02 | DISS |
| *Rnaset2b* | 2 | 9.60E-02 | DISS |
| *Rpsa* | 1.08 | 9.60E-02 | DISS |
| *Slc25a10* | 2.94 | 9.60E-02 | DISS |
| *Spry1* | 2.18 | 9.60E-02 | DISS |
| *Tango2* | 1.8 | 9.60E-02 | DISS |
| *Ubqln1* | -1.06 | 9.74E-02 | HOMO |
| *Gadd45b* | 1.49 | 9.83E-02 | DISS |
| *Atrx* | -1.06 | 9.92E-02 | HOMO |
| *Itpr3* | 3.12 | 9.92E-02 | DISS |
