## Supplementary material for "Hippocampal transcriptomic responses to cellular dissociation": Supp. Table 2

| Sanes & Lichtman Molecules | Related Transcripts |
| --- | --- |
| GLUTAMATE RECEPTORS | |
| GluR1; GluR2 | *Gria1; Gria2* |
| mGluR1; mGluR4; mGluR5; mGluR7 | *Grm1; Grm4; Grm5; Grm7* |
| NMDA NR2A; NMDA NR2D; NMDA NR1 | *Grin1; Grin2a; Grin2d* |
| OTHER NEUROTRANSMITTERS | |
| norepinephrine and b-adrenergic receptors | *Adrb1; Adrb2; Adrb3* |
| adenosine and adenosine 2A receptors | *Adra1a; Adra1b; Adra1d; Adra2a; Adra2b; Adra2c* |
| dopamine and D1 dopamine receptors | *Th; Drd1* |
| mu and delta opioid receptors | *Oprm1; Oprd1* |
| acetylcholine receptors | *Chrna1; Chrna7; Chrna3; Chrnb1; Chrnb2; Chrnb3* |
| muscarinic receptors | *Chrm1; Chrm2; Chrm3; Chrm4; Chrm5* |
| GABA receptors | *Gabra1; Gabra2; Gabra3; Gabra5; Gabra6* |
| GABA-B receptors | *Gabrb1; Gabrb2; Gabrb3* |
| cannabinoid receptor | *Cnr1; Cnr2* |
| orphanin NQ and nocioceptin receptors | *Pnoc; Oprl1;* |
| serotonin receptors | *Htr1a; Htr1b; Htr1f; Htr2a; Htr2c; Htr2b; Htr3a; Htr3b; Htr5a; Htr5b; Htr7; Htr6; Htr4;* |
| endothelin-1 | *Edn1* |
| gamma-aminobutyric acid (GHB) receptors | *Gabrr1; Gabbr1* |
| INTERCELLULAR MESSENGERS, THEIR SYNTHETIC ENZYMES AND THEIR RECEPTORS | |
| CO | *NA* |
| NO | *NA* |
| EGF | *Egf* |
| basic FGF | *Fgf2* |
| superoxide | *NA* |
| neuregulin | *Nrg1; Nrg2; Nrg3* |
| erbB4 | *Erbb4* |
| NGF | *Ngf* |
| BDNF | *Bdnf* |
| TrkB | *Ntrk2* |
| nNOS; eNOS | *Nos1; Nos3* |
| arachidonic acid | *NA* |
| platelet activating factor | *NA* |
| interleukin 1 beta | *Il1b* |
| H2S | *NA* |
| beta activin | *Inhba* |
| CALCIUM/CALMODULIN BINDING PROTEINS | |
| calmodulin | *Calm1; Calm2; Calm3* |
| RC3/neurogranin | *Nrgn* |
| calretinin | *Calb1; Calb2* |
| GAP43/B50/neuromodulin | *Gap43* |
| S100 | *S100b* |
| ION CHANNELS |  |
| L-type calcium channels | *Cacna1c; Cacna1d; Cacna1s; Cacna1f; Cacna1b; Cacna1a; Cacna1e* |
| olfactory cyclic nucleotide-gated channel | *Cnga2* |
| VESICLE- AND SYNAPSE-ASSOCIATED PROTEINS | |
| synaptophysin | *Syp* |
| a-SNAP | *Napa* |
| VAMP | *Vamp1; Vamp2; Vamp3; Vamp4; Vamp5; Vamp8* |
| rab3a | *Rab3a* |
| syntaxin 1B | *Stx1b;* |
| Synapsin I | *Syn1* |
| SNAP 25 | *Snap25* |
| PSD-95 | *Dlg4* |
| TRANSCRIPTION FACTORS | |
| Retinoic acid receptor beta | *Rarb* |
| CREB | *Creb1* |
| Krox 20; Krox 24 | *Egr1; Egr2* |
| ADHESION MOLECULES | |
| ephA5 | *Epha5* |
| ephrinA5 | *Efna5* |
| NCAM | *Ncam1; Ncam2* |
| E-cadherin; N-cadherin | *Cdh1; Cdh2* |
| thy-1 | *Thy1* |
| telencephalin | *Icam5* |
| L1/NgCAM | *L1cam* |
| HB-GAM/pleitrophin | *Ptn* |
| integrins | *Itga1; Itga10;Itga11; Itga2; Itga2b;Itga3; Itga4; Itga5; Itga6; Itga7; Itga8; Itga9; Itgad; Itgae; Itgal; Itgam; Itgav; Itgax; Itgb1; Itgb1bp1; Itgb2; Itgb2l; Itgb3; Itgb3bp; Itgb4; Itgb5; Itgb6; Itgb7; Itgb8; Itgbl1* |
| integrin-associated protein | *Cd47* |
| tenascin-C | *Tnc* |
| CARBOHYDRATES | |
| Polysialic acid | *NA* |
| Ganglioside GM1 | *NA* |
| Ganglioside GQ1B | *NA* |
| KINASES |  |
| inositol-triphosphate-3-kinase | *Itpka; Itpkb; Itpkc* |
| MAPK | *Mapk1; Mapk10; Mapk11; Mapk12; Mapk14; Mapk3; Mapk4; Mapk6; Mapk7; Mapk8; Mapk9* |
| src | *Src* |
| fyn | *Fyn* |
| protein kinase A C beta 1 subunit | *Prkacb* |
| protein kinase A RI beta subunit | *Prkar1b* |
| protein kinase C-gamma | *Prkcg* |
| protein kinase G | *Prkg1* |
| protein kinase M-zeta | *Prkcz* |
| CaM kinase I; II; IV | *Camk1; Camk2; Camk4* |
| ecto-protein kinase | *NA* |
| PROTEASES AND THEIR INHIBITORS | |
| calpain | *Capn1; Capn10; Capn11; Capn12; Capn13; Capn15; Capn2; Capn3; Capn5; Capn6; Capn7; Capn8; Capn9* |
| calpastatin | *Cast* |
| protease nexin 1 | *Serpine2* |
| tissue plasminogen activator | *Plat* |
| plasmin | *Plg* |
| E6-AP ubiquitin ligase | *Ube3a* |
| OTHER ENZYMES | |
| phospholipase A2 | *Pla2g10; Pla2g12a; Pla2g12b; Pla2g15; Pla2g16; Pla2g1b; Pla2g2a; Pla2g2c; Pla2g2d; Pla2g2e; Pla2g2f; Pla2g3; Pla2g4a; Pla2g4b; Pla2g4e; Pla2g4f; Pla2g5; Pla2g6; Pla2g7* |
| phospholipase C beta | *Plcb1; Plcb2; Plcb3; Plcb4* |
| phospholipase C gamma | *Plcg1; Plcg2* |
| ADP ribosyl transferase | *Parp1* |
| calcineurin | *Ppp3ca; Ppp3cb; Ppp3cc* |
| protein phosphatase I | *Phpt1* |
| acetylcholinesterase | *Ache* |
| adenylate cyclase | *Adcy1; Adcy10; Adcy2; Adcy3; Adcy4; Adcy5; Adcy6; Adcy7; Adcy8; Adcy9* |
| guanylate cyclase | *Gucy1a2; Gucy1a3; Gucy1b2; Gucy1b3; Gucy2c; Gucy2d; Gucy2e; Gucy2g* |
| MISCELLANEOUS | |
| Spectrin/fodrin | *Sptan1; Sptbn1* |
| GFAP | *Gfap* |
| Stathmin RB3/XB3 | *Stmn4* |
| EBI-1 G protein-coupled receptor | *Ccr7* |
| Mas G-protein coupled receptor | *Mas1* |
| Vesl | *Homer1; Homer2; Homer3* |
