## Supplemental Table 3 for "Hippocampal transcriptomic responses to cellular dissociation"

| marker | gene | lfc | padj | direction |
| --- | --- | --- | --- | --- |
| astrocyte | *ALDOC* | 0.93 | 3.48E-01 | neither |
| astrocyte | *AQP4* | -0.15 | 9.64E-01 | neither |
| astrocyte | *FGFR3* | 0.16 | 9.51E-01 | neither |
| astrocyte | *GFAP* | 0.71 | 6.15E-01 | neither |
| astrocyte | *GJB6* | 0.65 | 7.99E-01 | neither |
| astrocyte | *SLC1A2* | 0.04 | 9.87E-01 | neither |
| microglia | *CD68* | 2.35 | 9.11E-02 | DISS |
| microglia | *TNF* | 2.4 | 2.21E-02 | DISS |
| microglia | *PTPRC* | 1.37 | 7.28E-01 | neither |
| neuron | *GABRA1* | -1.05 | 1.41E-01 | neither |
| neuron | *KCNQ2* | -0.41 | 6.56E-01 | neither |
| neuron | *NEFH* | 0.59 | 7.47E-01 | neither |
| neuron | *NEFL* | 0.3 | 7.94E-01 | neither |
| neuron | *NEFM* | -0.37 | 7.10E-01 | neither |
| neuron | *SLC12A5* | -0.87 | 2.67E-01 | neither |
| neuron | *SNAP25* | 0.37 | 8.18E-01 | neither |
| neuron | *SV2B* | -0.07 | 9.95E-01 | neither |
| neuron | *SYT1* | -0.33 | 7.61E-01 | neither |
| oligodendrocyte | *GJC2* | 2.39 | 9.60E-02 | DISS |
| oligodendrocyte | *MAG* | 3.31 | 4.48E-05 | DISS |
| oligodendrocyte | *MAL* | 3.2 | 2.32E-04 | DISS |
| oligodendrocyte | *MBP* | 1.95 | 8.03E-03 | DISS |
| oligodendrocyte | *MOBP* | 2.6 | 4.41E-04 | DISS |
| oligodendrocyte | *MOG* | 2.48 | 2.27E-02 | DISS |
| oligodendrocyte | *SOX10* | 2.16 | 5.75E-02 | DISS |
| oligodendrocyte | *CSPG4* | 1.38 | 1.42E-01 | neither |
| oligodendrocyte | *GAL3ST1* | 2.1 | 5.14E-01 | neither |
| oligodendrocyte | *PDGFRA* | 1.24 | 2.39E-01 | neither |
| oligodendrocyte | *UGT8A* | 1.68 | 1.74E-01 | neither |
